# Cellular and molecular census of the pleiotropic disease *locus* Negr1 in zebrafish

**DOI:** 10.64898/2026.10.08.757516

**Authors:** Andrés González-Suarez, Laura Pola-Morell, Hernán López-Schier

## Abstract

A major challenge in precision medicine is to understand the cellular origin of disease-causing genetic alterations. One prominent example is the *Negr1 locus*, where common non-coding variants are associated with an unusually broad range of human traits and conditions, including body mass index (BMI), major depression, schizophrenia, autism-spectrum conditions and anorexia. Yet, no coding loss-of-function allele has been established as causal. These observations suggest that distinct *Negr1*-expressing cell populations underlie distinct traits. However, this hypothesis is difficult to test in tractable experimental systems without first determining where this gene may act. To overcome this limitation, we generated a knock-in reporter at the zebrafish *Negr1 locus* and combined single-cell transcriptomics with whole-brain registration and single-neuron projection mapping. We found that *Negr1* is expressed in a graded manner across neuronal classes of the brain and retina, radial glia and astrocyte-like cells, and most consistently in oligodendrocyte progenitor cells. Projection mapping suggests three conserved hubs formed by olfactory bulb projection neurons, tectal periventricular projection neurons and cerebellar eurydendroid cells. Our findings are consistent with the idea that *Negr1* pleiotropy arises from dose perturbation of a rather broadly expressed gene. More generally, this integrative approach to produce granular multimodal datasets should enable high-resolution etiological studies in models of human disease.

## INTRODUCTION

Disorders of the nervous system have a profoundly negative impact on the quality of life of affected individuals and their families. They are a major long-term burden to healthcare systems worldwide (Deuschl et al., 2020; GBD 2016 Neurology Collaborators, 2019; GBD 2017 US Neurological Disorders Collaborators, 2021; Kessler et al., 2003; Roehrig, 2016). Many neuropsychiatric traits have a genetic basis (Bray and O’Donovan, 2018; Geschwind and Flint, 2015). Consequently, historical efforts have been justifiably directed towards genome-wide association studies (GWAS) in well-characterized human cohorts, which have been enormously successful in revealing polymorphisms in the genome that correlate with illness. Yet, there are very few examples of disease-causing genetic variants for which the molecular mechanisms driving the association have been experimentally determined, hindering the development of targeted treatments (Gallagher and Chen-Plotkin, 2018; Song Y. et al., 2026). Moreover, a remarkable finding of GWAS is that most disease-associated variants lie in non-coding regions of the genome (Watanabe et al., 2019; Welter et al., 2014; Peng and Greally, 2026). A prominent example is the gene encoding Neuronal Growth Regulator 1 (NEGR1). Over 100 non-coding polymorphisms in the *NEGR1 locus* have been associated with various pathological traits in humans, including anorexia, obesity, autism-spectrum disorders, lower sociability, schizophrenia, major depression, learning difficulties and post-traumatic stress disorders (Figure 1) (Deng et al., 2022; Brugger et al., 2020; Rana and Mobin, 2020; Singh et al., 2019; Grove et al., 2019; Karis et al., 2018; Szczurkowska et al., 2018; Singh et al., 2018; Locke et al., 2015; Kim et al., 2014; Levey et al., 2021). However, the mechanistic link between variants in the *NEGR1 locus* and diseases that do not share obvious etiology remains obscure (Noh et al., 2020; Dall’Aglio et al., 2021). Disentangling *locus* pleiotropy from variant specificity is challenging because the cellular origin of these conditions is generally unknown. One way to understand how different genomic polymorphisms predispose an individual to a specific disease, and thus develop personalized treatment, is to identify and characterize the cells that express *NEGR1* (Williams et al., 2024). A novel approach towards this aim is to integrate genomic data from humans with experimental single-cell technologies in animal models (Haque et al., 2017; Shalek and Benson, 2017; Olislagers et al., 2022). The conserved anatomy and neurophysiology of zebrafish combined with a powerful genetic and behavioral framework makes it an ideal system to tackle this challenge (Kunst et al., 2019; Wee et al., 2019; Marques et al., 2018; Thyme et al., 2019; Espigares et al., 2022). Under this rationale, we generated a comprehensive high-resolution structural and molecular census of *negr1*-expressing cells in larval zebrafish.

**Figure 1.**
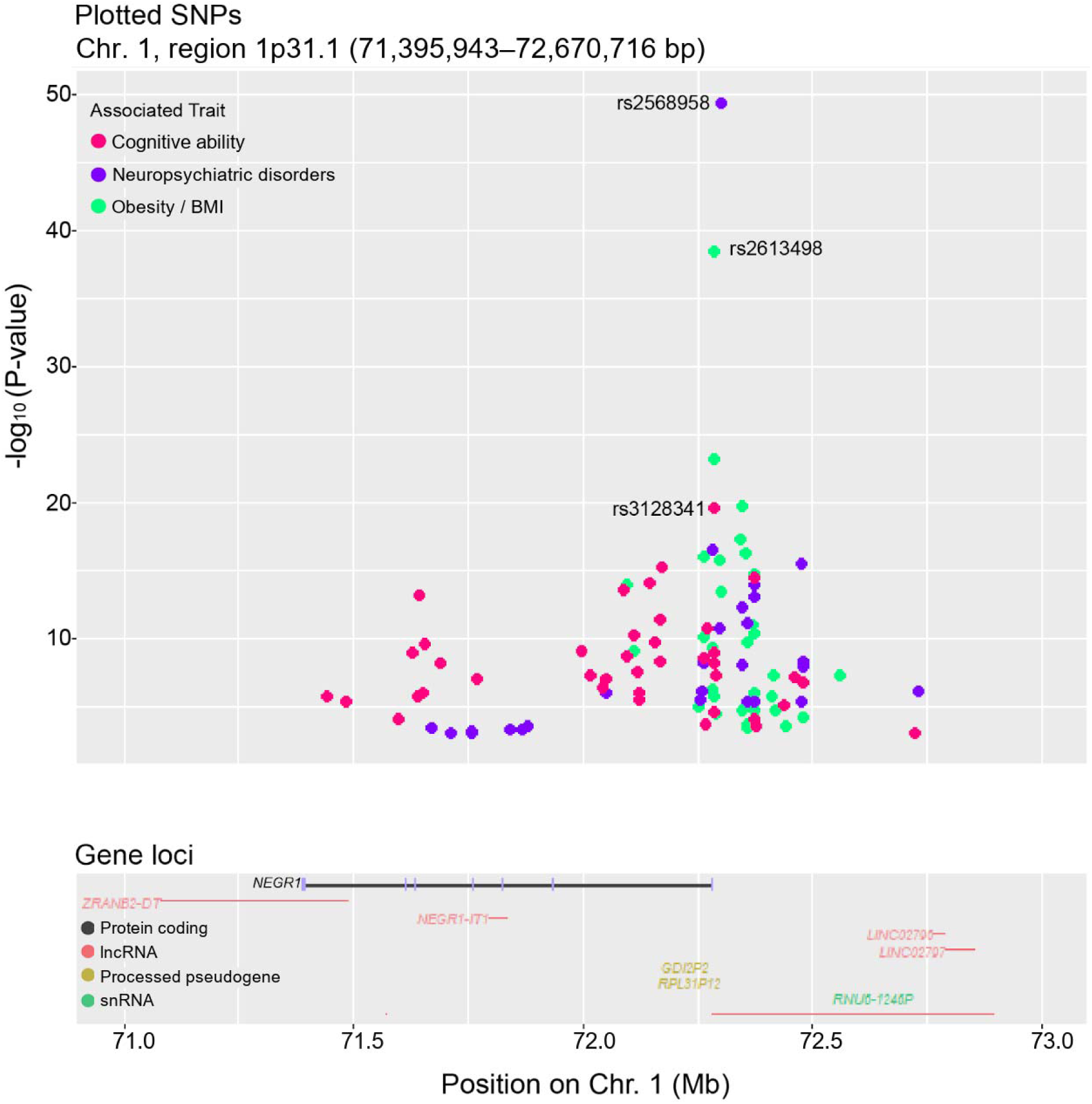
LocusZoom plot of *NEGR1* SNP associations with traits. The plot shows the genomic region of the human *NEGR1 locus* on chromosome 1. SNPs are shown as colored dots according to their association with a phenotypic trait within the *NEGR1 locus*. The top SNP for each trait is labeled with its reference SNP cluster ID (rsID). Genes in the region are shown below as horizontal color-coded lines indicating gene extent and biotype. *NEGR1* color code: introns in black and protein-coding exons in magenta. P values were −log10-transformed and extracted from different GWAS in which the *NEGR1* gene was associated with a trait.

## RESULTS

### Identification and characterization of *negr1* in zebrafish

NEGR1 belongs to the IgLON family of cell-adhesion proteins, which also includes the closely related NTM, LSAMP, OPCML (OBCAM) and IGLON5 (Venkannagari et al., 2020). Therefore, we used a stringent test stack to unambiguously identify the zebrafish ortholog of human *NEGR1*. We searched publicly available genomic data using BLAST and detected a candidate gene on zebrafish chromosome 6 with important diagnostic characteristics. First, its genomic structure is identical to that of the *bona fide Negr1* in mammals (Figure 2A). It contains 7 exons that encode a predicted protein of 346 amino acids, consisting of three Ig-like C2-type domains located at amino-acid positions 33-123, 140-241 and 226-313. The same domains are found in the human protein at equivalent positions 38-134, 139-221 and 225-313, and in the murine counterpart within corresponding positions 32-128, 133-215 and 219-307. A signal peptide and a C-terminal GPI-anchor signal are respectively encoded by exons 1 and 7, both in the mammalian and zebrafish proteins. Furthermore, using the Reciprocal Best Hit Pipeline, we identified conserved local synteny of human chromosome 1 and zebrafish chromosome 6, as well as in the orthologous chromosomes of mouse and macaque (Figure 2B) (Catchen et al., 2009). Finally, the genomic sequence revealed the presence of homologous neighboring genes in human and zebrafish (Figure 2C). Together, these data identify it as the zebrafish ortholog of *NEGR1*, hereafter *negr1*. Consistent with this, independent data from Ensembl Compara annotates *negr1* (ENSDARG00000021607) as the single one-to-one zebrafish ortholog of human *NEGR1*, indicating that a possible paralog arising from the teleost-specific genome duplication was not retained (Dyer et al., 2025).

**Figure 2.**
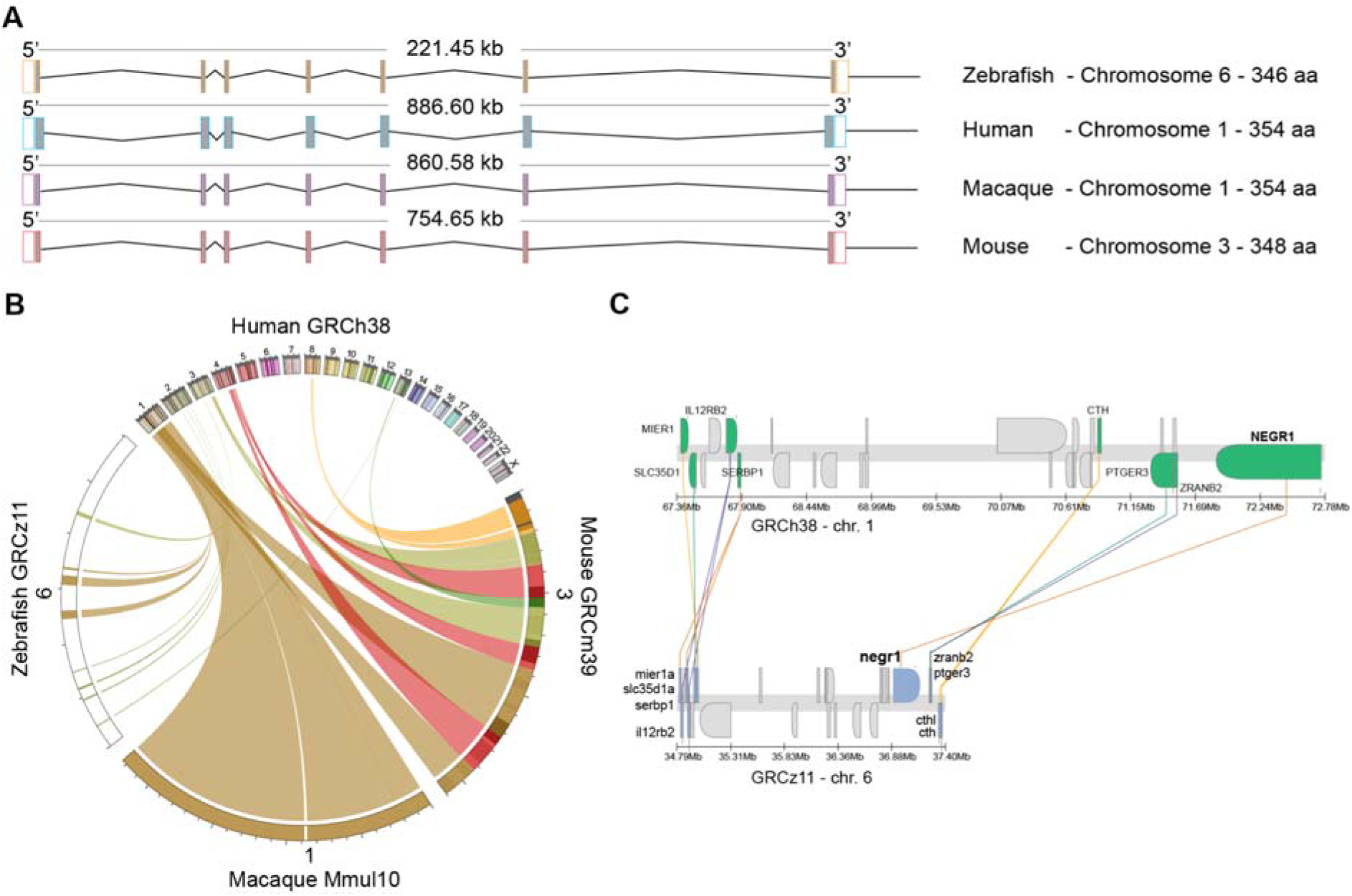
Identification of *negr1* in zebrafish. (A) Alignment of the *NEGR1* gene structure indicating the number of exons, gene length, chromosome (Chr) location and number of amino acids in different species. (B) Circos plot of chromosomal relationships between human (22 autosomes and Chr X) and macaque (Chr 1), mouse (Chr 3) and zebrafish (Chr 6). Chromosomes are arranged in a circle; ribbons connect genomic regions that correspond to positions of orthologous genes between species. (C) Schematic of syntenic conservation between human (Chr 1) and zebrafish (Chr 6), highlighting the conservation of the *NEGR1 locus* between these species.

### Reporter line for *negr1* in zebrafish

Immunocytological studies have localized NEGR1 in the membranes of mammalian neurons (Funatsu et al., 1999), with no *in vivo* data. There is no known antibody that specifically recognizes Negr1 in zebrafish. Therefore, to reveal zebrafish Negr1 localization *in vivo*, we fused the full-length *negr1* cDNA to EGFP and cloned the fusion into a vector containing 10 linked repeats of the Upstream Activation Sequence (UAS) to express it using a GAL4 driver. A resulting stable transgenic line, Tg(UAS:*negr1*-EGFP), was crossed with fish carrying HuC:GAL4 to express Negr1-EGFP in neurons of the central nervous system (Figure 3A-C). As expected, we detected a strong signal at the plasma membrane and a weaker but homogeneous signal in the cytosol, with signal excluding from the nucleus (Figure 3C). Next, we used CRISPR/Cas9-mediated genome engineering to knock-in an EGFP reporter at the *negr1 locus* (Kimura et al., 2014). Of note, this insertion neither replaces nor tags the endogenous gene, and is therefore not expected to alter *negr1* expression or function (Figure 3D). We obtained germline transmission of a strongly fluorescent line, Tg(*negr1*:EGFP^hzm17^), whose spatiotemporal expression *in vivo* was characterized in embryonic and larval specimens (Figure 3E-H). We detected the first signal at approximately 20 hours post fertilization (hpf) (data not shown) in small bilaterally symmetric regions of the brain. At 2 days post fertilization (dpf) expression increases in the optic tectum, the midbrain and hindbrain (Figure 3E), with high expression in the cerebellum and the medulla oblongata. We also saw consistent but low expression in the forebrain. Between 4 dpf and 6 dpf, we detected *negr1*:EGFP in the telencephalon and in part of the olfactory bulb and spinal cord. Next, we compared the patterns of *negr1*:EGFP and the endogenous *negr1* transcript at 6 dpf by whole-mount fluorescent *in situ* hybridization (HCR RNA-FISH) (Figure 3F-K). The two signals largely overlapped. Therefore, we decided to conduct further work with 6 dpf animals because this stage conveniently enables cross-platform integration of our data with community-generated light- and electron-microscopy brain atlases, whole-brain activity mapping and behavioral studies (Hildebrand et al., 2017; Thyme et al., 2019; Shainer et al., 2023; Zada et al., 2024; Shang et al., 2024).

**Figure 3.**
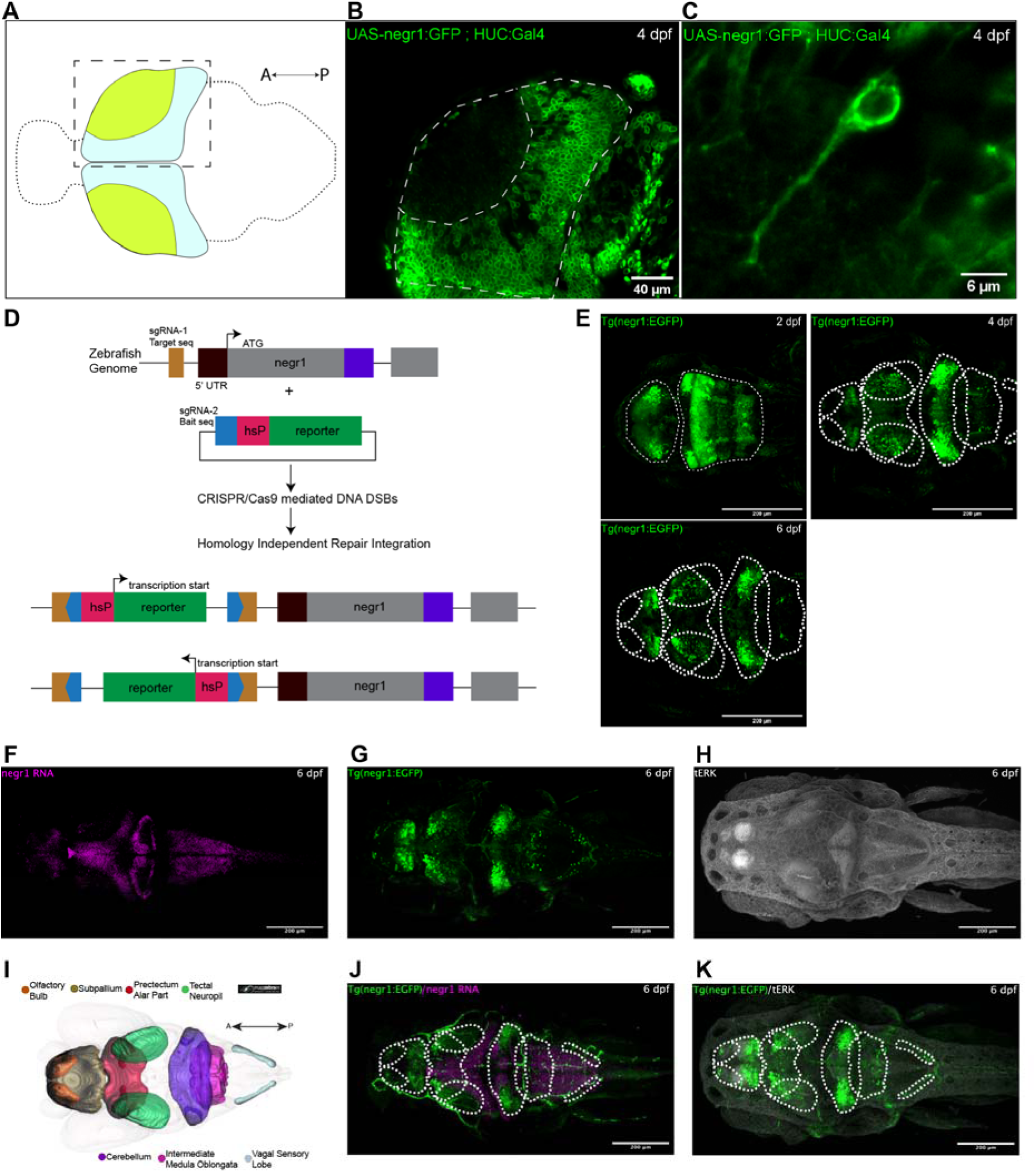
Anatomical characterization of *negr1* expression in larval zebrafish. Laser-scanning single- and and two-photon confocal images (dorsal views) of double transgenic Tg(UAS:*negr1*-EGFP; HuC:GAL4) and Tg(*negr1*:EGFP) zebrafish. (A) Schematic representation of the zebrafish brain (dorsal view); the dashed box marks the optic tectum region of the right hemisphere shown in B. (B) Confocal image (dorsal view) of the right tectal hemisphere of a 4 dpf double transgenic larva. (C) Close-up of a tectal neuron (right hemisphere) expressing Negr1-EGFP at the cell membrane and along its axonal projection. (D) Simplified scheme of the CRISPR/Cas9-mediated Knock-in approach used to generate the knock-in transgenic line (adapted from Kimura et al., 2014). (E) Two-photon images of Tg(*negr1*:EGFP) from 2 dpf to 6 dpf. Maximum-intensity projections show clusters of EGFP signal from *negr1*-expressing cells in the larval brain at different developmental stages. The expression remains bilaterally symmetric over time. Dotted lines highlight brain regions with localized clusters of *negr1*-expressing cells. (F-H) Whole-mount fluorescence *in situ* hybridization and immunohistochemistry in 6 dpf larvae. Images are maximum-intensity projections of confocal z-stacks. (F) HCR RNA-FISH for *negr1*. (G) EGFP signal from *negr1*-expressing cells in the larval brain. (H) tERK staining. (I) 3D brain template of a 6 dpf zebrafish showing, color-coded by region, the brain regions in which *negr1*-expressing cells are localized: forebrain (FB: olfactory bulb and subpallium), midbrain (MB: pretectum alar part and tectal neuropil) and hindbrain (HB: cerebellum, intermediate medulla oblongata and vagal sensory lobe). Registration was done using the multimodal atlas of the larval zebrafish brain (MapZebrain, Baier lab). (J-K) Maximum-intensity projections of confocal z-stacks of 6 dpf Tg(*negr1*:EGFP) knock-in larvae. (J) Composite image of Tg(*negr1*:EGFP) (green) with *negr1* HCR RNA-FISH (magenta). (K) Composite image of Tg(*negr1*:EGFP) with tERK staining. Anterior is to the left and posterior to the right. Dotted lines highlight brain regions with localized clusters of *negr1*-expressing cells.

### Anatomical organization of *negr1*-expressing cells

To place *negr1*-positive cells in a mesoscopic anatomical frame, we stained *negr1*:EGFP animals with an antibody against total ERK (tERK) (Figure 3H, K), which facilitates registration onto the Max Planck Zebrafish Brain atlas (MapZebrain) (https://mapzebrain.org) (Kunst et al., 2019; Shainer et al., 2023; Schulze et al., 2023). MapZebrain is a consolidated reference that standardizes many transgenic reporter lines in well-defined brain regions of larval zebrafish and, importantly, also contains annotated projections of over 4,000 traced individual neurons. We captured images from both sagittal and horizontal planes, and the resulting raw data were processed and rendered three-dimensional using Fiji and 3D Viewer (Supp. Video 1) (Schmid et al., 2010). Analysis of the registration revealed strong *negr1*:EGFP expression in the forebrain, the olfactory bulb (OB) and the subpallium (Sp); within the hindbrain, the most robust expression was localized to the cerebellum (Ce) (Bae et al., 2009) (Figure 3I-K). We also localized *negr1*:EGFP in parts of the spinal cord, the midbrain, tectal neuropil (TN), the pretectal alar plate (Pre), the intermediate medulla oblongata (MO) and vagal sensory lobe (VL) (Figure 3I-K).

### Molecular classification of *negr1*-positive cells

Because the main aim of this study is to molecularly identify the cells that express *negr1* at single-cell resolution, we isolated green-fluorescent cells from approximately 180 heads of 6 dpf *negr1*:EGFP transgenic larvae and processed them along a standard pipeline for single-cell RNA sequencing (scRNA-seq) (Figure 4A) (Scott et al., 2021; Cosacak et al., 2019). We sequenced 9,195 individual cells using the 10x Genomics Chromium platform. This dataset was then analyzed using Scanpy (Wolf et al., 2018; Luecken and Theis, 2019). After doublet removal and quality control, 7,363 cells were retained, with a median of 1,749 genes detected per cell. Principal component analysis (PCA) and Leiden clustering (see Methods) resolved 23 cell clusters (clusters 0-22; Figure 4B), which we annotated using literature-curated diagnostic transcripts and unbiased marker-gene analysis (Supp. Figure 2; Supp. Tables 1-2). Because whole heads were dissociated, the dataset contains cells from the brain, retina, inner ear and craniofacial tissues. The largest group comprised craniofacial mesenchymal derivatives, consistent with a cranial neural crest origin (Howard et al., 2021; Schilling and Kimmel, 1994; Raible et al., 1992): mesenchyme and fibroblasts (cluster 0; *col1a1a*, *col1a2*, *dlx5a*), chondrocytes (cluster 2; *col2a1b*, *matn4*), fibroblast-like mesenchyme (cluster 3) and smooth-muscle-like cells (cluster 4; *rbpms2b*, *acta2*). Other non-neural clusters included epithelial cells (cluster 13; *epcam*, *cdh1*), *six1b*+ epithelium (cluster 10), inner-ear (otic) epithelium (cluster 17; *otomp*, *stm*) (Söllner et al., 2003; Murayama et al., 2005), vascular endothelial cells (cluster 15; *fgd5a*, *kdrl*), muscle-like cells (cluster 21; *fbp2*, *eno3*) and a small population of proximal tubule-like cells (cluster 22; *agxtb*, *si:ch211-139a5.9*). Retinal clusters comprised cycling retinal progenitors (cluster 6; *rx1*, *vsx2*, *atoh7*, *mki67*), retinal neurogenic precursors (cluster 8), Müller glia and late retinal progenitors (cluster 11; *rx1*, *vsx2*, *slc6a9*), cone photoreceptors (cluster 7; *crx*, *nr2e3*, *tulp1a*, *opn1lw2*) and their cycling precursors (cluster 14), bipolar cells (cluster 12; *vsx1*, *gabrr2a*, *calb2b*) and amacrine cells (cluster 20; *tfap2a*, *tfap2b*, *pax6b*, *slc32a1*). No cluster expressed the retinal ganglion cell markers *isl2b* or *pou4f1*. Clusters from the brain comprised a mixed population of differentiating and mature neurons (cluster 5; *elavl3*, *elavl4*, *nrxn3a*, *neurod1*), radial glia (cluster 1; *her4.1*, *her15.1*, *sox2*, *fabp7a*), cycling radial glia and neural progenitors (cluster 9; *her4.1*, *her15.1*, *mki67*, *top2a*), astrocyte-like radial glia (cluster 18; *glula*, *fabp7a*, *slc1a2b*, *s100b*) (Lam et al., 2009; Bernardos and Raymond, 2006; Lange et al., 2020), oligodendrocyte progenitor cells (OPCs; cluster 16; *olig1*, *olig2*, *sox10*, *cspg4*, *nkx2.2a*) (Xiao et al., 2022; Marisca et al., 2020; Mangin and Gallo, 2011; Bradl and Lassmann, 2010; Park et al., 2007; El Waly et al., 2014; Tan et al., 2014; Othman et al., 2011) and floor plate cells (cluster 19; *shha*, *foxa2*, *slit1b*, *ntn1b*, *spon1b*). The floor plate cluster also expressed *pkd1b* and *urp2*, which mark cerebrospinal fluid-contacting neurons (Quan et al., 2015), but lacked neuronal genes such as *elavl3* and *snap25a* (Supp. Figure 2).

**Figure 4.**
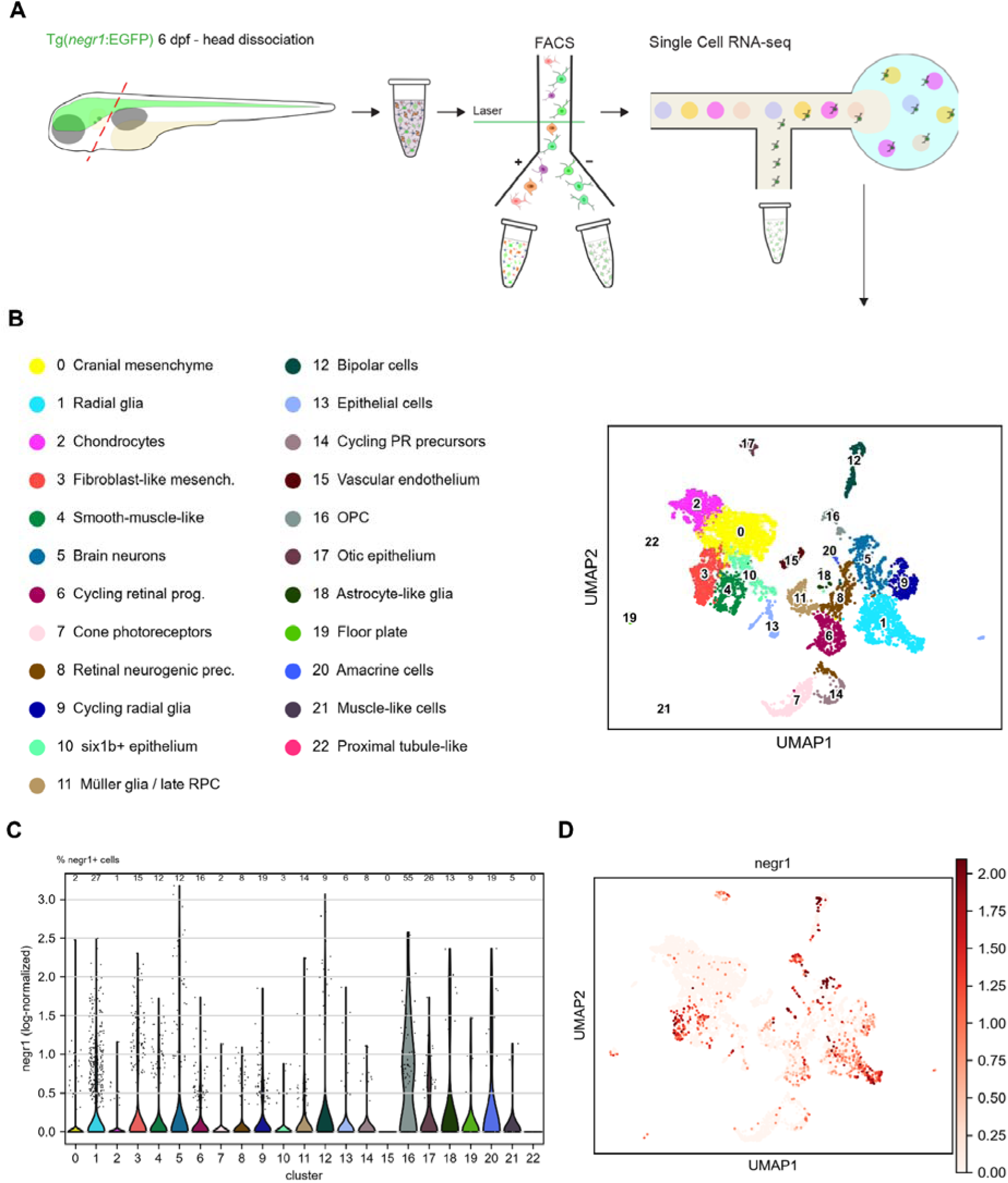
Molecular profiling of *negr1*-expressing cells. (A) Experimental workflow for the dissociation, isolation and scRNA-seq of *negr1*-expressing cells. Heads from 6 dpf Tg(*negr1*:EGFP) larvae were dissociated to obtain a single-cell suspension, followed by FACS to isolate GFP-positive cells and droplet encapsulation on the 10x Chromium platform for library generation. (B) Uniform manifold approximation and projection (UMAP) of 7,363 cells arranged in 23 clusters (0-22), with their annotation (left). Clusters were generated with the unsupervised Leiden algorithm and are arranged in two dimensions by transcriptional similarity. (C) Violin plot of *negr1* expression in each cluster; numbers above the violins indicate the percentage of *negr1*-positive cells. *negr1* is detected in all clusters except clusters 15 and 22, and the highest fraction of positive cells is found in OPCs (cluster 16). (D) UMAP visualization of *negr1* expression. Color intensity represents the level of expression (log-normalized counts).

Using the above data, we next assessed the spatial distribution of selected cell classes in the brain. Using double-transgenic Tg(*negr1*:EGFP);Tg(*olig1*:mScarlet) larvae, we found sparse but consistent co-localization of both markers in the hindbrain and the spinal cord (Supp. Figure 1A, a1 - a3). We also performed hybridization chain reaction RNA fluorescence *in situ* hybridization (HCR RNA-FISH) in *negr1*:EGFP transgenics with a set of custom probes. *hoxb5a*, as a marker for motor neurons (Barsh et al., 2017), was found in a cluster of *negr1*+ cells in the posterior part of the hindbrain (Supp. Figure 1B, b1 - b3), suggesting motor-neuron identity. *olig2* is a marker for eurydendroid cells (ENs) (Supp. Table 1 and Supp. Figure 1C) (Harmon et al., 2020). We found co-expression of *olig2* and *negr1* in symmetric clusters in the cerebellum (Supp. Figure 1C, c1 - c3). Finally, we found co-expression of *negr1* and *id2b*, a marker of tectal pyramidal neurons (TLPN; DeMarco et al., 2020; DeMarco et al., 2021), in neurons deep within the tectal neuropil layer of the optic tectum (OT) (Supp. Figure 1D, d1 - d3).

### *negr1* spans glial and neuronal clusters

We next examined how *negr1* expression is distributed across neuronal and glial identities (Figures 4C, 4D and 5). *negr1* transcripts were detected in 21 of the 23 clusters, but the fraction of positive cells varied widely, from below 2% in chondrocytes, mesenchyme and cone photoreceptors to 55% in OPCs (Figure 4C). Pan-neuronal and synaptic genes (*elavl3*, *elavl4*, *snap25a*, *snap25b*, *sv2a*, *sypb* and *syt1a*) identified brain neurons (cluster 5), cone photoreceptors (cluster 7), bipolar cells (cluster 12) and amacrine cells (cluster 20) as mature neuronal populations (Figure 5A, C). Of these, *negr1* was detected in 19% of amacrine cells, 12% of brain neurons and 9% of bipolar cells, but in only 2% of cone photoreceptors. Among glia, *negr1* was detected in 27% of radial glia (cluster 1), 19% of cycling radial glia and neural progenitors (cluster 9) and 13% of astrocyte-like radial glia (cluster 18) (Figure 5B, C). Both radial glia clusters expressed the Notch target genes *her4.1*, *her4.2* and *her15.1*, but whereas most cells of cluster 1 lacked proliferation markers, cluster 9 was strongly proliferative (*mki67* and *top2a* in nearly all cells), indicating that *negr1* is expressed in both non-cycling and proliferating radial glia. OPCs (cluster 16) were the only population in which most cells expressed *negr1* (Figure 4C). Most OPCs also expressed synaptic and neuronal genes (*elavl3*, *snap25a*, *sv2a* and *syt1a*) and AMPA and NMDA receptor subunits (75-88% of cells; Figures 5A and 6B). This profile was shared by all OPC subclusters and was not associated with elevated doublet scores. The co-expression of *olig1*/*olig2* with synaptic and glutamate receptor genes could indicate progenitors transitioning to a neuronal fate; however, it is more parsimoniously explained by OPC identity, as OPCs receive glutamatergic synapses (Bergles et al., 2000; Marisca et al., 2020).

**Figure 5.**
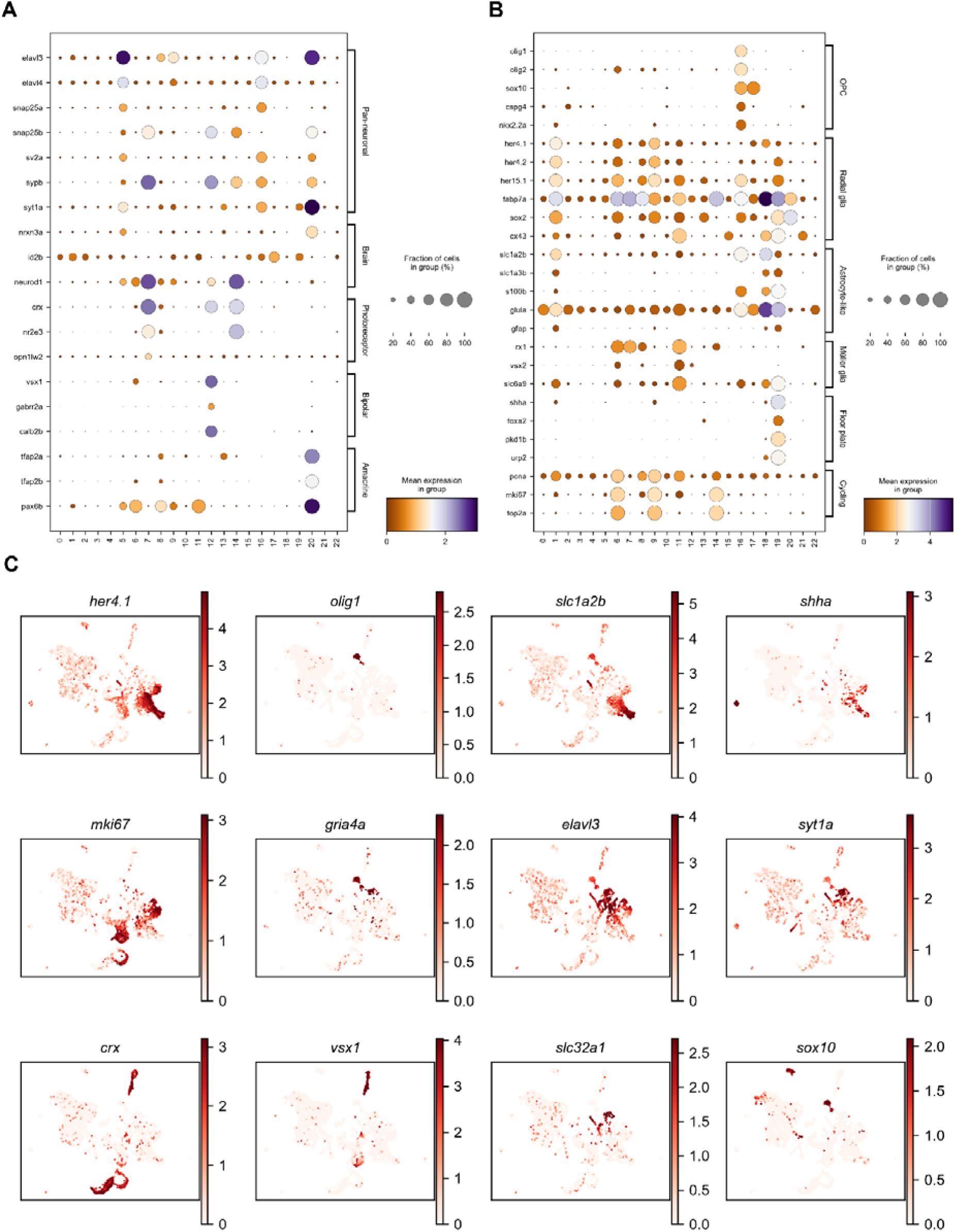
Neuronal and glial gene markers. (A) Dot plot of pan-neuronal and synaptic genes and of markers of brain neurons, photoreceptors, bipolar and amacrine cells. (B) Dot plot of markers of OPCs, radial glia, astrocyte-like glia, Müller glia, floor plate and proliferating cells. Dot size indicates the fraction of cells expressing each gene and color the mean expression in each cluster. (C) UMAP plots of selected glial (*her4.1*, *olig1*, *slc1a2b* and *sox10*), floor plate (*shha*), proliferation (*mki67*), glutamate receptor (*gria4a*), neuronal (*elavl3* and *syt1a*), retinal (*crx* and *vsx1*) and GABAergic (*slc32a1*) genes. Color intensity is proportional to the expression level of each gene.

**Figure 6.**
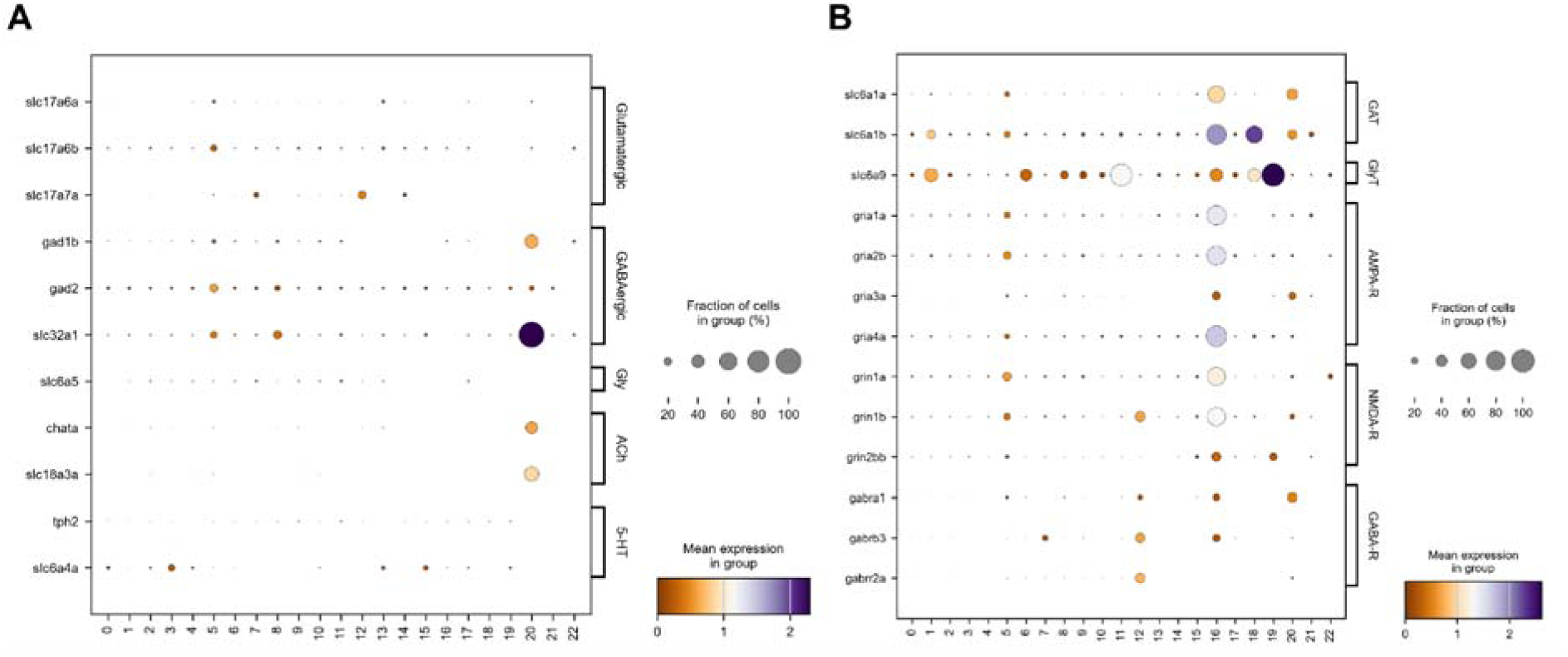
Neurotransmitter profiles of *negr1*-expressing clusters. (A) Genes for neurotransmitter synthesis and vesicular transport defining glutamatergic, GABAergic, glycinergic (Gly), cholinergic (ACh) and serotonergic (5-HT) identity. Clear GABAergic and cholinergic signatures are restricted to amacrine cells (cluster 20), whereas glycinergic and serotonergic markers are essentially absent. (B) GABA (GAT) and glycine (GlyT) transporters and AMPA, NMDA and GABA-A receptor (R) subunits. Transporters are expressed mainly by glial clusters, and glutamate receptor subunits by OPCs (cluster 16). Dot size indicates the fraction of expressing cells and color the mean expression.

### Neurotransmitter profile of *negr1*-expressing neuronal clusters

We next characterized the neurotransmitter identity of the annotated clusters using genes for transmitter synthesis and vesicular transport (Figure 6A). Amacrine cells (cluster 20) showed a clear GABAergic (*slc32a1* in all cells, *gad1b*) and cholinergic (*chata*, *slc18a3a*) signature, consistent with starburst amacrine cells. Brain neurons (cluster 5) contained both GABAergic (*gad2*, *slc32a1*) and glutamatergic (*slc17a6b*) cells, and the vesicular glutamate transporter gene *slc17a7a* was expressed in bipolar cells and photoreceptors. Markers of glycinergic (*slc6a5*) and serotonergic (*tph2*) neurons were essentially absent. The GABA transporter genes *slc6a1a* and *slc6a1b* and the glycine transporter gene *slc6a9* were expressed mainly by glia, including OPCs, astrocyte-like radial glia, floor plate and Müller glia (Figure 6B), consistent with neurotransmitter uptake by glial cells. GABAergic neurons are abundant in the larval zebrafish optic tectum (Robles et al., 2011; Barker et al., 2021), a region of strong *negr1*:EGFP expression (Figure 3). Notably, NEGR1 has also been linked to inhibitory neurons in the mammalian brain, where it co-localizes with parvalbumin-positive interneurons in human brain tissue (Muthuraman et al., 2026), its loss reduces the number of parvalbumin-positive interneurons in the mouse hippocampus (Singh et al., 2019), and it promotes the synaptic targeting of the GABA-synthesizing enzyme GAD65 (Su et al., 2025). Similarly, we characterized the neuropeptide and neurotrophic-factor profile of *negr1*-expressing cells by taking the human gene groups Neuropeptides (#1902), Neurotrophins (#1585) and GDNF family ligands (#1586) from the HUGO Gene Nomenclature Committee (HGNC) database, identified their zebrafish orthologs, added their receptors, and examined their expression across all clusters (Figure 7). Neuropeptide genes were generally sparse. The most prominent were the secretogranin gene *scg2b* and the chromogranin gene *chga*, both expressed by most OPCs, *adcyap1b* in brain neurons and OPCs, *apln* in cone photoreceptors, and *igf2b* in mesenchymal and endothelial cells (Figure 7A). Among receptors, the apelin receptor genes *aplnra* and *aplnrb* were highly expressed in OPCs and in cycling photoreceptor precursors; apelin signaling has been implicated in many physiological processes (Zhang et al., 2018). The insulin and IGF1 receptor genes *insra*, *insrb* and *igf1rb* were broadly expressed at intermediate levels, in line with the role of insulin signaling in zebrafish neurogenesis (Gence et al., 2023). Neurotrophins regulate the development, maintenance and function of the vertebrate nervous system (Huang and Reichardt, 2001). Neurotrophin receptor genes were enriched in OPCs, which expressed *ngfra* (53% of cells), *ntrk3a* (49%) and *ntrk3b* (70%). *ngfra* was also expressed in amacrine cells (Figure 7B). The nerve growth factor receptor (NGFR) regulates the growth and differentiation of sympathetic and sensory neurons (Micera et al., 2007). GDNF-family ligands and receptors were sparse, with *gfra1a* in amacrine cells and *ret* and *gfra3* in epithelial cells (Figure 7C). In zebrafish, *gdnf* regulates the differentiation of diencephalic dopaminergic neurons (Wong et al., 2021), is required for enteric nervous system development (Shepherd et al., 2001), and guides developing and regenerating lateral line axons (Schuster et al., 2010).

**Figure 7.**
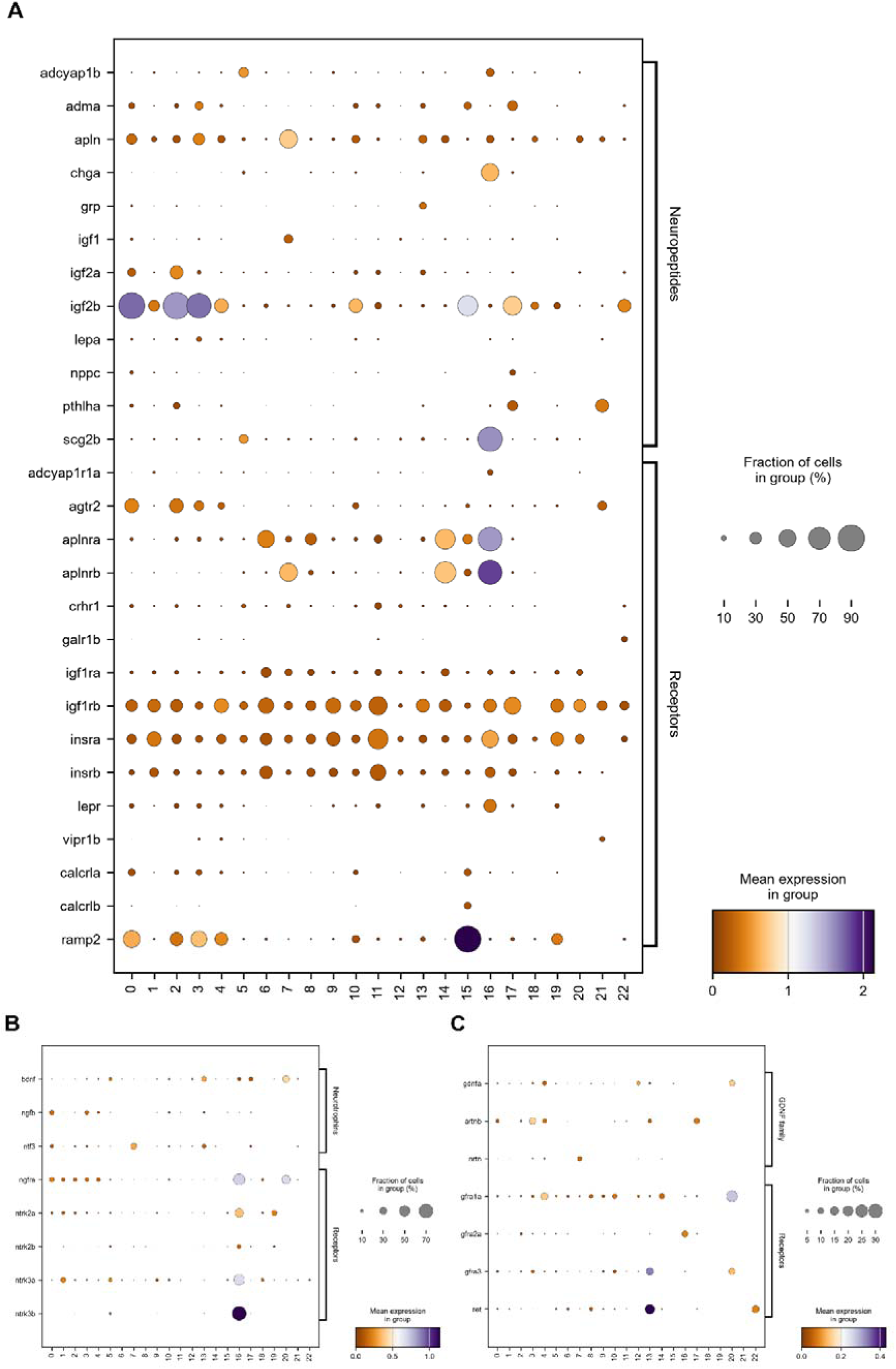
Neuropeptides, receptors and neurotrophic factors in *negr1*-expressing cells. (A) Zebrafish orthologs of the HGNC Neuropeptides gene group (#1902) and their receptors; only genes detected in at least 10% of cells in at least one cluster are shown. (B) Neurotrophins (HGNC #1585) and their receptors, which are enriched in OPCs (cluster 16). (C) GDNF-family ligands (HGNC #1586) and their receptors. Dot size indicates the fraction of expressing cells and color the mean expression in each cluster.

### Topographic organization of *negr1*-expressing cells

We combined our anatomical and molecular data to infer the projection patterns of *negr1*-expressing cells at single-cell resolution. Because individual neuronal morphologies cannot be resolved directly in our reporter line, we registered *negr1*:EGFP to the MapZebrain atlas and identified catalogued single neurons whose somata lie within *negr1*:EGFP-positive regions (see Methods). We focused on the three populations best represented in our dataset: olfactory bulb projection neurons (OBpn), periventricular projection neurons (PVPN) and eurydendroid cells (ENs). In the olfactory bulb (OB), *negr1*:EGFP-positive territory overlapped with the somata of OBpn located in its most anterior-dorsal part. These neurons project to five brain areas: the posterior telencephalon (pTL), ventral telencephalon or subpallium (vTL-Sp), habenula (Hb), posterior tuberculum (PT) and the OB itself (Figure 8A) (Miyasaka et al., 2014), suggesting that *negr1*-expressing OB neurons may share this projection pattern. In the optic tectum, PVPN with somata in *negr1*:EGFP-positive regions project across the midbrain-hindbrain boundary to the most anterior part of the hindbrain and the medulla oblongata (MO) (Figure 8B). In the cerebellum, ENs located within *negr1*:EGFP-positive regions project to parts of the optic tectum and hindbrain, most of them targeting the pretectum and the tectal neuropil (Figure 8C). Orthogonal views of the registered data refined these patterns: OBpn project mainly to the subpallium (vTL-Sp) (Figure 8D); PVPN have a more distributed output, partially targeting the deep ventral diencephalon and mainly the MO (Figure 8E); and ENs show the broadest distribution, extending to the hypothalamus, tegmentum, thalamus, oculomotor nucleus and MO (Figure 8F). These three populations therefore do not form a single circuit but three separate long-range hubs, each projecting to multiple downstream regions.

**Figure 8.**
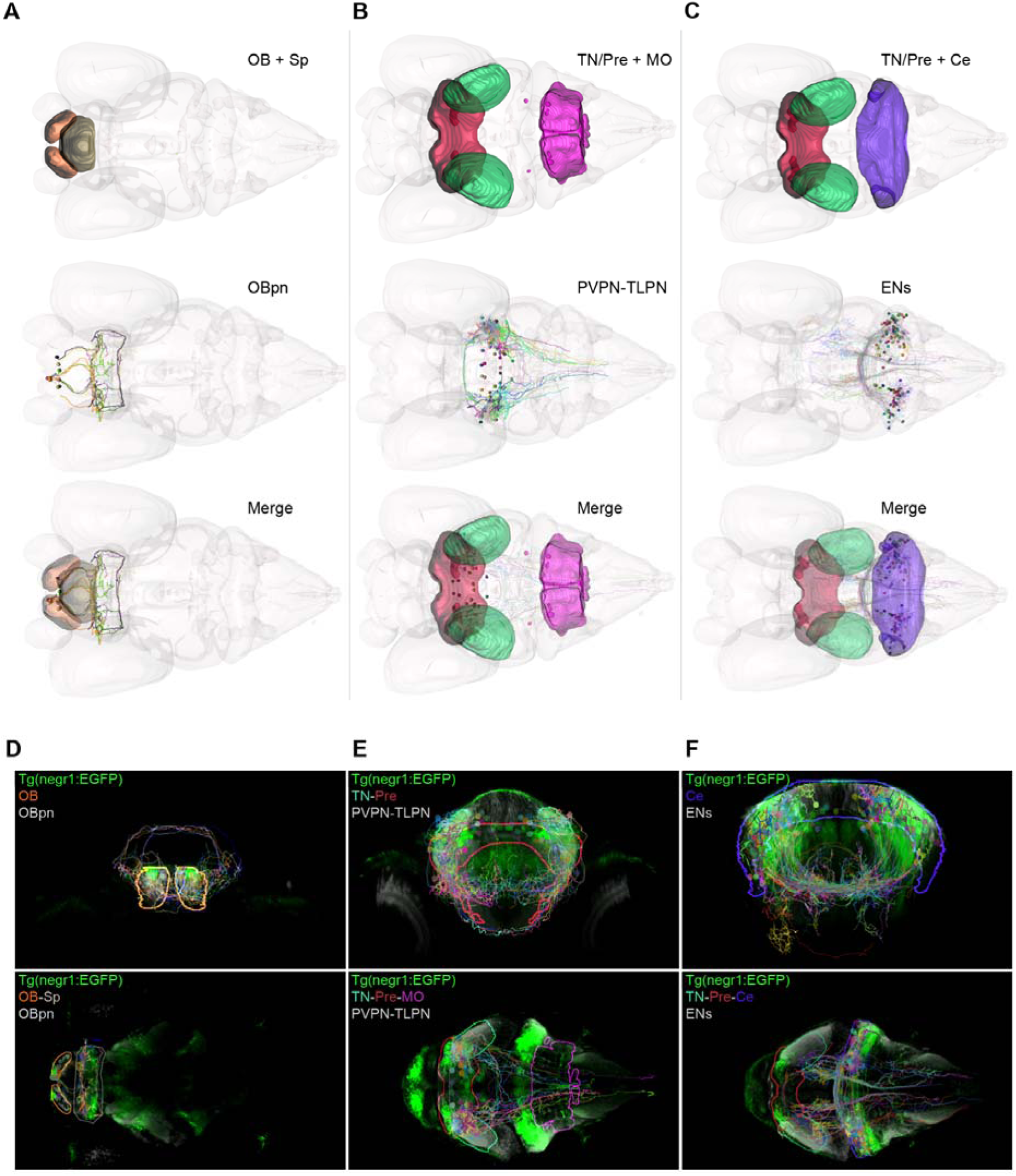
Topographic organization of *negr1*-expressing cells. (A-C) 3D renderings of the standardized 6 dpf zebrafish brain showing manually defined regions and single-neuron projections located within distinct brain areas. (D-F) Single orthogonal views (coronal and horizontal) of Tg(*negr1*:EGFP) registered to the atlas. Coronal views show neurons located in different brain regions and their projections. Horizontal views show neuronal somata and their projections targeting distinct brain regions. Abbreviations: OB+Sp (olfactory bulb and subpallium), OBpn (olfactory bulb projection neurons), TN/Pre+MO (tectal neuropil/pretectum alar part and intermediate medulla oblongata), PVPN-TLPN (periventricular projection neurons - tectal pyramidal neurons), Ce (cerebellum) and ENs (eurydendroid cells). Horizontal views: anterior to the left, posterior to the right. Coronal views: dorsal up, ventral down.

## DISCUSSION

According to the World Health Organization, nearly 1.2 billion people worldwide, equivalent to 1 in 7 individuals, live with a mental health condition (World Health Organization, 2026), many of which arise from genetic predisposition. The genomic variants that contribute to them are overwhelmingly non-coding, individually small in effect, and shared across separate diagnoses (Watanabe et al., 2019; Cross-Disorder Group of the Psychiatric Genomics Consortium, 2019). *NEGR1* shows all three of the above properties and presents a paradox. Common non-coding variation at 1p31.1, where the *NEGR1 locus* is located, is associated with body mass index, major depression, schizophrenia, autism-spectrum traits, cognitive performance and anorexia. Yet, no recurrent coding loss-of-function allele has ever been established as causal for any of them, and animal models have not converged on a consensus conclusion. For example, ablation of *Negr1* in mice produces a modest reduction in body mass (Lee et al., 2012), whereas gene knock-down in mice and rats increases body weight and food intake (Boender et al., 2014). Behavioral and structural phenotypes across studies remain incomplete (Singh et al., 2018; Singh et al., 2019; Szczurkowska et al., 2018). The most likely explanation is that a gene with such a pleiotropic profile is unlikely to act through a single cell type. Therefore, determining where it acts, at cellular resolution and in an intact vertebrate brain, is a necessary step toward interpreting the associations between genomic alterations and multiple disjointed diseases.

As a first step towards this goal, we produced a molecular and cellular census in zebrafish, which reveals that *negr1* is detected in 21 transcriptionally defined populations in the larval head, albeit with a fraction of positive cells ranging from less than 2% to 55%. Together with the reporter line that we generted, validated by endogenous transcript staining, and HCR RNA-FISH, our data show that *negr1* is expressed in several neuronal classes, including brain neurons, retinal amacrine and bipolar cells, cerebellar eurydendroid cells and tectal pyramidal neurons. It is also expressed in non-cycling and proliferating radial glia, in astrocyte-like radial glia, in oligodendrocyte progenitors and, at lower levels, in mesenchymal, epithelial and inner-ear cells. We found that *negr1* expression is graded rather than binary, most consistently expressed in oligodendrocyte progenitor cells (cluster 16), whereas the highest individual expression levels occur in subsets of brain neurons and bipolar cells (clusters 5 and 12), and other glial and neuronal clusters show lower, more heterogeneous expression (Figure 4C). These data suggest that *locus*-associated traits arise not from loss of Negr1 function in a trait-specific cell type, but instead from dose perturbation of a broadly deployed molecule. Under this scenario, non-coding variants that alter *NEGR1* expression levels would modify many neuronal circuits only weakly, rather than perturbing one circuit strongly. This would produce persistent associations of small effect size distributed across different physiological systems. Supporting this conclusion, *NEGR1* expression is elevated in the prefrontal cortex of individuals with major depressive disorder relative to controls (Wang et al., 2020; Li et al., 2020), and overexpression of *Negr1* in the mouse prefrontal cortex and hippocampus is sufficient to induce anxiety- and depression-like phenotypes (Zhang et al., 2025). Moreover, partial loss experiments have yielded phenotypes opposite to those of null mutant alleles (Boender et al., 2014; Lee et al., 2012). Therefore, the most parsimonious explanation is that *NEGR1*-associated traits reflect dosage sensitivity across a graded expression landscape in different cell types, rather than complete loss of function. This further suggests that dosage sensitivity of an unevenly distributed gene across cell types requires buffering. The IgLON family, to which NEGR1 belongs, includes LSAMP, NTM, OPCML and IGLON5. These proteins assemble as homo- and hetero-dimers with partially overlapping binding preferences (Venkannagari et al., 2020). Complete loss of one member may therefore be compensated within cells that co-express a redundant partner, while dosage shifts in cells expressing *NEGR1* in isolation may not be buffered. This may explain why the complete ablation of *Negr1* in rodents has proved a poor model of the human-associated conditions. Some evidence supports this explanation (El-Brolosy et al., 2019), which warrants testing the hypothesis in future studies.

Moreover, if any perturbation is weak and distributed, its phenotypic consequences should converge on multiple cell types rather than on single cell identities. By registering *negr1*:EGFP to the MapZebrain reference and identifying catalogued single neurons within *negr1*-positive regions, we identified three putative long-range hubs. Olfactory bulb projection neurons target the subpallium, posterior telencephalon, habenula and posterior tuberculum. Tectal periventricular projection neurons project across the midbrain-hindbrain boundary to the medulla oblongata. Cerebellar eurydendroid cells distribute broadly to hypothalamus, thalamus, tegmentum, oculomotor nucleus and medulla. These are conserved long-range pathways, whose mammalian counterparts underlie olfactory-limbic connections, sensorimotor action and hypothalamic control of feeding. These pathways map onto the trait domains associated with the *NEGR1 locus* in human GWAS. We propose these units as candidates for trait vulnerability. This foundational work offers a path to interpreting non-coding pleiotropic *loci* combining anatomical, molecular and physiological experimentation.

### Limitations of the study

Single biological studies cannot offer a definitive answer, and this one is no exception. First, all analyses were performed at a single larval stage (6 dpf) and therefore do not capture how *negr1* expression changes later in development or in the adult brain. Second, the scRNA-seq dataset derives from a single pooled sample of FACS-sorted head cells, which includes non-neural tissues such as the retina and inner ear, and lacks biological replicates. In addition, *negr1* transcripts were detected in only 11% of the sorted cells, as expected for a low-abundance transcript in droplet-based scRNA-seq, so the absence of detection in individual cells or small clusters should not be interpreted as absence of expression. Third, the knock-in reporter reports promoter activity rather than endogenous transcript or protein levels; although HCR RNA-FISH confirmed a largely overlapping pattern, the two may differ in some cell types. In addition, the genomic integration site and orientation of the knock-in allele were not confirmed molecularly, so the identity of the line rests on germline-transmitted EGFP expression that reproduces the endogenous *negr1* transcript pattern (Figure 3F, J). Fourth, the projection maps rely on neurons whose somata lie within *negr1*:EGFP-positive regions, rather than on neurons directly confirmed to express *negr1*. Finally, we did not manipulate *negr1* dosage, so the dosage model we propose remains a hypothesis to be tested in future loss-of-function and dosage experiments.

## Materials and Methods

### Ethics Statement

All experiments were performed according to the guidelines by the Ethical Committee of Animal Experimentation of the Helmholtz Zentrum München, protocol number ROB-55.2-2532.Vet_02-21-117 approved by the “Regierung von Oberbayern”, Germany. Experiments with wild-type and transgenic zebrafish were done in specimens of undetermined sex up to 7 dpf.

### Zebrafish Lines and Husbandry

Zebrafish eggs and larvae were kept under standard conditions at 28.5°C in 0.3× Danieau’s solution, with a 14 h:10 h light:dark cycle. The following lines were newly generated for this study: Tg(*negr1*:EGFP^hzm17^) and Tg(UAS:*negr1*-EGFP). We used the following previously described lines: Tg(*olig1*:mScarlet) (Xiao et al., 2022), and Tg(HuC:GAL4) (Faucherre and López-Schier, 2011).

### Generation of Tg(*negr1*:EGFP) and Tg(UAS:*negr1*-EGFP) Transgenic Zebrafish

The Tg(*negr1*:EGFP) line was generated using the knock-in strategy of Kimura et al. (2014). One-cell-stage wild-type embryos were injected with the Mbait-hsp70:EGFP donor plasmid, Cas9 mRNA (Sigma), the Mbait sgRNA, which linearizes the donor *in vivo*, and two sgRNAs targeting the *negr1* promoter region (sgRNA-1 and sgRNA-2; Supp. Table 4). The two genomic target sites lie 180 and 428 bp upstream of the transcription start site, so that neither the 5′ UTR nor the coding sequence is disrupted. Carrier fish were identified in the following generation by EGFP fluorescence. sgRNA templates were produced by annealing a gene-specific oligonucleotide carrying a T7 promoter to a constant oligonucleotide encoding the sgRNA scaffold (Gagnon et al., 2014), followed by *in vitro* transcription. For the Tg(UAS:*negr1*-EGFP) line, the DNA construct UAS:*negr1*-EGFP and transposase mRNA were injected as previously described (Kawakami, 2004). The UAS:EGFP DNA construct was generated using the Tol2kit entry vector (Kwan et al., 2007); the EGFP coding sequence was then fused downstream of the zebrafish *negr1* coding sequence, resulting in the final expression construct UAS:*negr1*-EGFP.

### Whole Mount HCR Fluorescence *in situ* Hybridization

HCR reagents, including probes, hairpins and buffers, were custom-designed and purchased from Molecular Instruments (Los Angeles, California, USA). Probe sets for *negr1*, *hoxb5a*, *olig2* and *id2b* each consisted of 20 split-initiator probe pairs per target. The staining was performed according to the protocol “*In situ* v3.0 protocol for whole-mount zebrafish larvae” (Choi et al., 2018). Tg(*negr1*:EGFP) larvae were grown until 6 dpf under standard conditions, anesthetized with MS-222 (tricaine, 0.16 g/L) and fixed with ice-cold 4% paraformaldehyde (PFA) overnight at 4°C with gentle shaking. Following fixation, 6-8 larvae were transferred into a 1.5 ml Eppendorf tube, washed in PBST (PBS + 0.1% Tween-20) for 3 × 5 min, dehydrated with 100% MeOH for 4 × 10 min and cooled to -20°C. Rehydration was performed in a MeOH/PBST series (5 min each: 100% MeOH; 75% MeOH: 25% PBST; 50% MeOH: 50% PBST; 25% MeOH: 75% PBST; 100% PBST), followed by 4 × 5 min washes in PBST. Permeabilization was carried out at room temperature in 30 µg/ml Proteinase K for 45 min, followed by two quick washes with PBST. A post-fixation with 4% PFA for 20 min was followed by 5 × 5 min washes in PBST. Probe hybridization buffer was pre-warmed at 37°C (manufacturer’s protocol), and larvae were pre-hybridized for 30 min; subsequently, the samples were incubated in the probe solution (2 µl of 1 µM probe stock in 500 µl probe hybridization buffer) overnight at 37°C. After removing the probe solution, samples were washed 4 × 15 min with probe wash buffer at 37°C and incubated in amplification buffer. After pre-amplification, the samples were incubated in the hairpin mixture (10 µl of 3 µM stock + 500 µl amplification buffer) overnight in the dark at room temperature. Lastly, amplified samples were washed 2 × 5 min, 2 × 30 min and 1 × 5 min in 5× SSCT. Following HCR RNA-FISH, larvae were stained with DAPI (4’,6-diamidino-2-phenylindole, 1:500, Sigma) at room temperature, followed by 3 × 10 min washes in PBST. Samples were either mounted for imaging or stored in 30% glycerol in PBST in the dark at 4°C.

### Immunohistochemistry

Whole-mount immunostaining was carried out as follows. First, zebrafish larvae at 2-6 dpf were fixed in ice-cold 4% PFA in PBST (0.1% Tween-20) overnight at 4°C. Second, samples were washed 3 × 10 min with PBST at room temperature, incubated for 2 h at room temperature in blocking solution (10% BSA in PBST) and subsequently incubated in primary antibodies overnight at 4°C. Third, samples were washed in PBST for 2 h, changing to fresh PBST every 30 min, then incubated with secondary antibodies at 4°C overnight. Antibodies and dilutions: mouse anti-MAPK-Erk, 1:500 (Cell Signaling, 4696); donkey anti-mouse Alexa Fluor® 594, donkey anti-mouse Alexa Fluor® 488. After incubation, samples were washed 3 × 5 min with PBST and stained with DAPI (as described for HCR RNA-FISH).

### Image Acquisition

Tg(*negr1*:EGFP) larvae at 2-6 dpf were imaged after fluorescence *in situ* hybridization (FISH) or *in vivo* for colocalization with Tg(*olig1*:mScarlet). Samples were embedded in 1% low-melting point agarose. For *in vivo* imaging, larvae were anesthetized in MS-222 (tricaine) 0.16 g/L. Images were acquired with a Zeiss LSM 880 Airyscan confocal laser scanning microscope (upright system) equipped with a 20× water immersion objective (W Plan-Apochromat 20×/1.0 DIC, WD 1.7 mm, Zeiss). Images of Tg(*negr1*:EGFP) at different developmental stages were acquired using a two-photon microscope (Sutter MOM) with a 16× water immersion objective (16×/0.80 NA W, WD 3.0 mm, Nikon). All images and z-stacks (2 tiles) were processed with Fiji (Schindelin et al., 2012), with contrast adjusted separately for each channel.

### Sample Preparation for scRNAseq

#### Tissue Dissociation

The protocol for larval zebrafish brain tissue dissociation was adapted from “Protocol for Dissection and Dissociation of Zebrafish Telencephalon for Single-Cell Sequencing” (Cosacak et al., 2020). Approximately 180 zebrafish larvae at 6 dpf were anesthetized (as described above) and immersed in ice-cold Ringer’s solution (without Ca^2+^ and Mg^2+^, 0.5 mM EDTA). Heads were carefully cut with a surgical scalpel blade (No. 10, Swann-Morton Ltd) at the boundary between hindbrain and spinal cord. Heads were collected into a 15 ml Falcon tube filled with fresh ice-cold Ringer’s solution (without Ca^2+^ and Mg^2+^, 0.5 mM EDTA added) and placed on ice. Collected heads were washed three times without incubation with 1× DPBS-EDTA (Dulbecco’s PBS, 1 mM EDTA) at room temperature, followed by incubation for 1 hour at 28°C with Buffer mix I and enzyme as described in the manufacturer’s protocol (Neural Tissue Dissociation Kit (P), Miltenyi Biotec), with mechanical dissociation by glass pipette every 10 min. The dissociated tissue solution was collected and filtered twice through a 30 µm strainer to remove cell aggregates. Subsequently, cells were resuspended in sort buffer (5% BSA, 1 mM EDTA in PBS) and kept on ice prior to cell sorting.

#### Fluorescence Activated Cell Sorting

After dissociation, the single-cell suspension was stained with 7-AAD (5 µg/mL, BD) and immediately FAC-sorted. GFP-positive cells from Tg(*negr1*:EGFP) were detected on a FACSAria™ (BD Biosciences) after 488 nm excitation; cells were passed through a 75 µm nozzle and collected in sort collection buffer (Ringer’s solution without Ca^2^+, 2% FBS, 10 µM EDTA). Only cells sorted in the first 30 min were used, to avoid sequencing stressed cells. GFP+ gates were set against non-fluorescent wild-type cells.

#### Single Cell RNA sequencing 10x Chromium

Single-cell RNA sequencing was performed using the 10x Chromium Single Cell 3’ Library & Gel Bead Kit v2 (PN-120237). Sorted cells were loaded into the 10x Chromium chip as described in the manufacturer’s protocol. Cells were lysed during encapsulation within the droplet, and polyadenylated RNA bound to the barcoded beads. The droplets were then reverse-transcribed, and the cDNA was purified and quality-checked. The libraries were sequenced to a depth of approximately 70,000 reads per cell on a HiSeq 4000 (Illumina). Raw data were pre-processed with Cell Ranger (v2.2.0, 10x Genomics). Data were demultiplexed and mapped to the zebrafish reference genome GRCz11 to generate single cell feature counts.

#### 10x Chromium Data Processing and Quality Control by Scanpy

Expression matrices were analyzed with the Python package Scanpy (v1.12.4) (Wolf et al., 2018) in Python 3.12; the complete software environment is provided with the analysis code. Doublets were first identified on raw counts with Scrublet (see Doublet Extraction). Standard quality control (QC) covariates were then computed: the number of counts per barcode, the number of genes per barcode and the fractions of counts from mitochondrial, ribosomal and hemoglobin genes per barcode (Ilicic et al., 2016; Griffiths et al., 2018). A high fraction of mitochondrial genes, few detected genes and low count depth are indicative of cellular stress or of cells whose cytoplasmic mRNA has leaked out (Luecken and Theis, 2019). We excluded cells with >20% mitochondrial counts, <3% ribosomal counts, >20% hemoglobin counts, >40,000 total counts, or fewer than 500 or more than 4,250 detected genes, and removed genes detected in fewer than 25 cells. This resulted in 7,363 cells and 16,735 genes. Counts were normalized to 10,000 per cell and log-transformed, and the effect of total counts per cell was regressed out. The top 5,000 highly variable genes (cell_ranger flavor) were used for principal component analysis (PCA). Next, we computed the neighborhood graph (15 neighbors, 40 principal components) and clustered the data using the Leiden algorithm (resolution 0.4; random seed 0). Cell clusters were visualized using a 2-dimensional UMAP (uniform manifold approximation and projection). Finally, to assign a molecular cell identity to each cluster, we identified marker genes with the Wilcoxon rank-sum test (Benjamini-Hochberg-adjusted P < 0.05, log fold change > 0.5) and annotated clusters manually using these markers and literature-curated diagnostic genes (Supp. Figure 2; Supp. Tables 1-2). For the neuropeptide analysis, the HGNC gene groups Neuropeptides (#1902), Neurotrophins (#1585) and GDNF family ligands (#1586) were mapped to their zebrafish orthologs, and genes detected in at least 10% of cells of at least one cluster were plotted together with their receptors.

#### Doublet Extraction

A common limitation of droplet-based scRNA-seq is the formation of technical artifacts, droplets that contain two cells, known as “doublets” (McGinnis et al., 2019). To overcome this issue, we applied Scrublet (v0.2.3) (Wolock et al., 2019) to the raw count matrix before quality control. Cells with a doublet score of 0.40 or higher were defined as doublets and removed (9,195 to 9,100 cells).

#### Curated SNPs and plotting

In this study, over 100 genome-wide association studies (GWAS) were reviewed. For each study, the inclusion criterion was an association with a trait or disease within the *NEGR1 locus*. To this end, we used Open Targets Genetics (Ghoussaini et al., 2021), which integrates genetic associations curated from the literature (GWAS) as well as newly derived loci from UK Biobank and FinnGen, and allows identification and prioritization of potential therapeutic drug targets. SNPs associated with *NEGR1* were obtained using an overall association score (0-1) (https://platform-docs.opentargets.org/associations) and were grouped into three main trait categories: obesity, cognitive ability and neuropsychiatric disorders (see Supp. Table 3). Subsequently, each curated SNP was plotted in a Locus Zoom-style plot using its −log10(P) value and aligned to its genomic position on chromosome 1. Plotting was done in R (RStudio) using the genome assembly GRCh38 as reference.

## Data and Code Availability

The original scRNA-seq dataset generated in this study is available at the National Center for Biotechnology Information Gene Expression Omnibus (GEO) under accession number GSE213794. The code for the Scanpy data analysis (https://github.com/sbohmgu/negr1_scRNAseq).

## Acknowledgements

This work was supported by the NIH BRAIN Initiative grant 1U19NS104653-01 (U.S.A.) and the NYUAD (U.A.E.) to HL-S. We thank Dr. Eva Laurell for help with MapZebrain.

## Notes

https://github.com/sbohmgu/negr1_scRNAseq

### Competing Interest Statement

H.L-S. is a scientific advisor for the biotechnology company Sensorion. The company had no role in this study. No conflict of interest exists.

