## Supplemental Figures for "Cellular and molecular census of the pleiotropic disease *locus* Negr1 in zebrafish"

**Supplementary Information**

- 3D rendering video

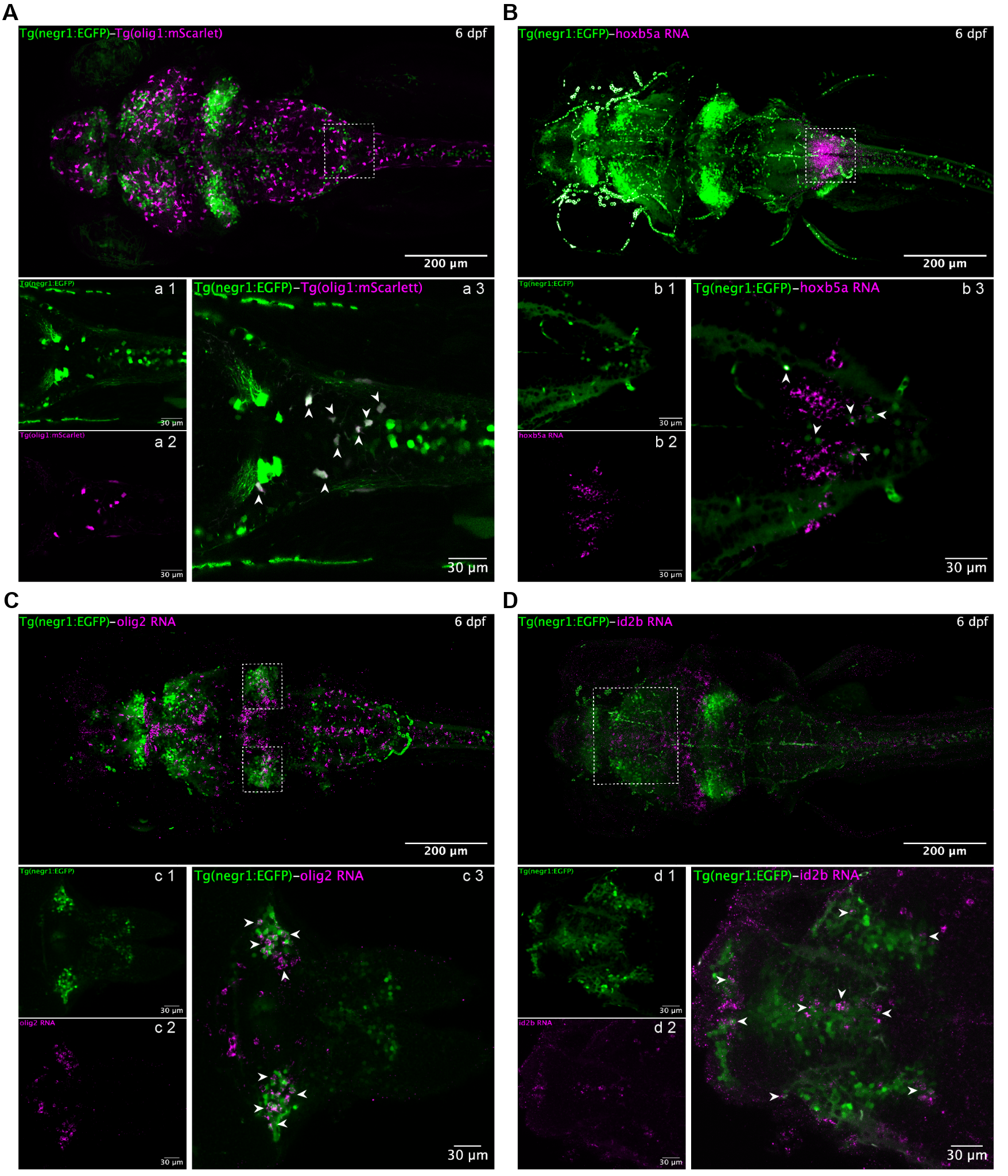

**Supp. Figure 1. Validation of neural cell types *in vivo*.** (A) *In vivo* confocal composite image (dorsal view) of a 6 dpf double transgenic larva: Tg(*negr1*:EGFP) in green and Tg(*olig1*:mScarlet) in magenta. The dashed box marks a region at the hindbrain-spinal cord (HB-SC) boundary. (a1-a2) Magnified single channels of the boxed region: *negr1*-expressing cells (a1, green) and *olig1*-expressing oligodendrocyte progenitor cells (OPCs) (a2, magenta). (a3) Merged image of the same region; white arrowheads indicate co-expression of *negr1*:EGFP and *olig1*:mScarlet in OPCs. (B-D) Whole-mount HCR RNA-FISH in 6 dpf Tg(*negr1*:EGFP) larvae reveals co-expression of marker genes within specific brain regions (dashed boxes). (b1-d1) Green channel: *negr1*-expressing cells. (b2-d2) Magenta channel: RNA probes for *hoxb5a* (motor neurons), *olig2* (eurydendroid cells) and *id2b* (periventricular neurons - tectal pyramidal neurons). (b3-d3) Merged images showing co-expression. Arrowheads indicate cells in which marker genes are co-expressed with *negr1*:EGFP. Anterior to the left, posterior to the right.

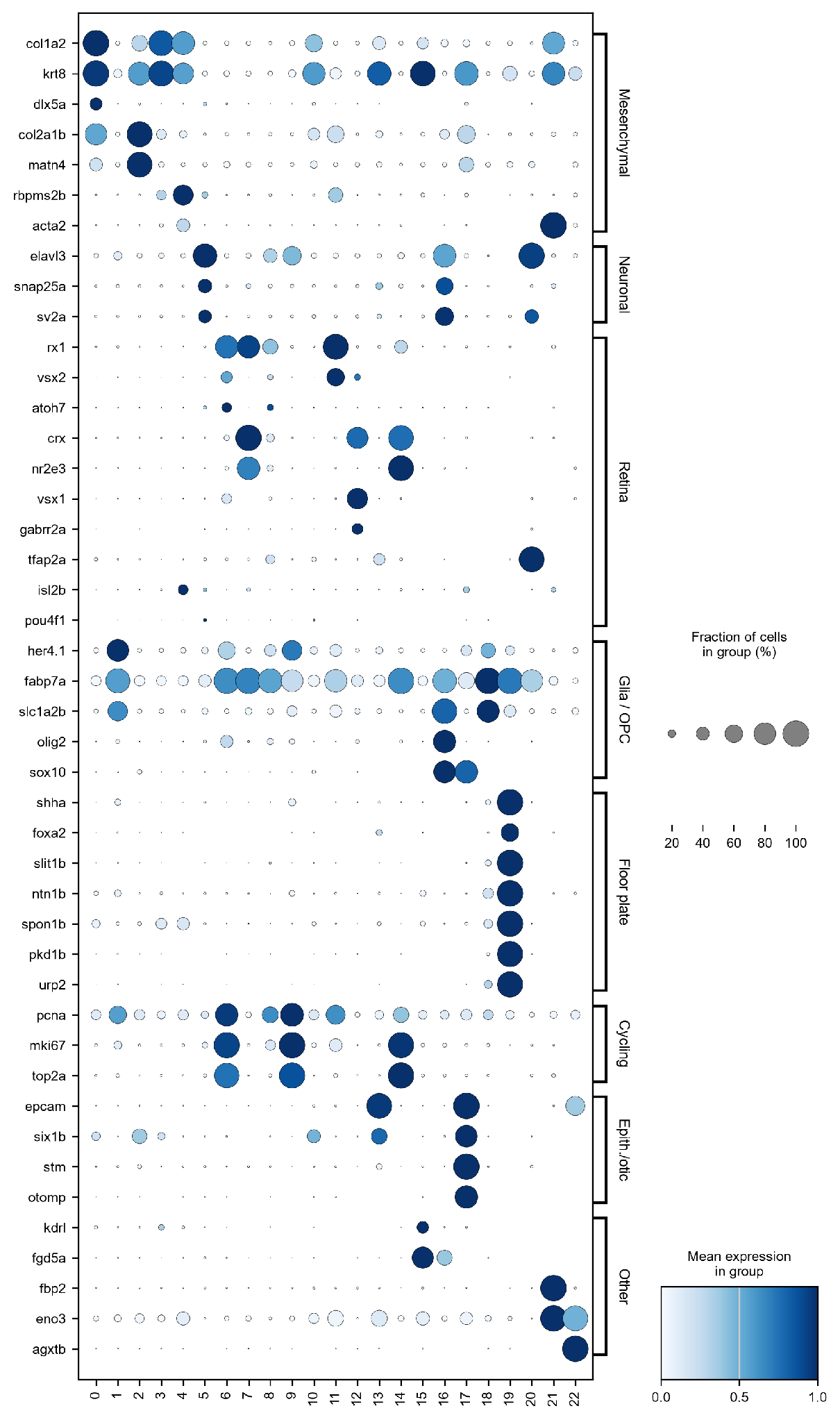

**Supp. Figure 2. Marker genes supporting cluster annotation.** Dot plot of diagnostic genes for each of the 23 clusters (color: mean expression scaled per gene; dot size: fraction of expressing cells). Cluster 19 expresses floor plate genes (*shha*, *foxa2*, *slit1b*, *ntn1b* and *spon1b*) together with *pkd1b* and *urp2*, but not the pan-neuronal genes *elavl3*, *snap25a* or *sv2a*. No cluster expresses the retinal ganglion cell markers *isl2b* or *pou4f1*.

**Supp. Table 4. Oligonucleotides used to generate Tg(*negr1*:EGFP).** Genomic coordinates refer to the GRCz11 assembly; the transcription start site of *negr1* is chr6:36,942,966 and the ATG is at chr6:36,943,152.

| **Name** | **Spacer (5′→3′)** | **PAM** | **Strand** | **Target site (GRCz11)** |
| --- | --- | --- | --- | --- |
| sgRNA-1 | GGTTGCGGGGCGATAAACCC | CGG | + | chr6:36,942,769–36,942,788 |
| sgRNA-2 | GGGCTTCAACTGTCGGTCTG | TGG | − | chr6:36,942,536–36,942,555 |
| Mbait | GGCTGCTGCGGTTCCAGAGG | — | — | donor plasmid (not genomic) |

| **Oligonucleotide** | **Sequence (5′→3′)** |
| --- | --- |
| KI-negr1-sgRNA_1 | GGGTTTATCGCCCCGCAACCCTATAGTGAGTCGTATTACGC |
| KI-negr1-sgRNA_2 | CAGACCGACAGTTGAAGCCCCTATAGTGAGTCGTATTACGC |
| KI-negr1-sgRNA_Mbait | CCTCTGGAACCGCAGCAGCCCTATAGTGAGTCGTATTACGC |
| Constant (scaffold) oligonucleotide | AAAAGCACCGACTCGGTGCCACTTTTTCAAGTTGATAACGGACTAGCCTTATTTTAACTTGCTATTTCTAGCTCTAAAAC |

Gene-specific oligonucleotides are shown in the antisense orientation and carry the reverse complement of the T7 promoter at their 3′ end; each was annealed to the constant oligonucleotide and transcribed *in vitro* (Gagnon et al., 2014). The Mbait sgRNA does not target the zebrafish genome and serves to linearize the donor plasmid.
